# Transcranial alternating current stimulation does not modulate corticospinal activity in humans

**DOI:** 10.1101/2022.01.13.476093

**Authors:** J. Ibáñez, B. Zicher, K. Brown, L. Rocchi, A. Casolo, A. Del Vecchio, D. Spampinato, C-A. Vollette, J. C. Rothwell, S. N. Baker, D. Farina

## Abstract

Transcranial alternating current stimulation (TACS) is commonly used to synchronise the output of a cortical area to other parts of the nervous system, but evidence for this based on brain recordings in humans is challenging. The brain transmits beta oscillations (^~^21Hz) to tonically contracted limb muscles linearly and through the fastest corticospinal pathways. Therefore, muscle activity may be used as a proxy measure for the level of beta entrainment in the corticospinal tract due to TACS over motor cortex. Here, we assessed if TACS is able to modulate the neural inputs to muscles, which would provide an indirect evidence for TACS-driven neural entrainment. In the first part of this study, we ran a series of simulations of motor neuron (MN) pools receiving inputs from corticospinal neurons with different levels of beta entrainment. Results indicated that MNs should be highly sensitive to changes in corticospinal beta activity. Then, we ran experiments on healthy human subjects (N=10) in which TACS (at 1mA) was delivered over the motor cortex at 21Hz (beta stimulation), or at 7Hz or 40Hz (control conditions) while the abductor digiti minimi (ADM) or the tibialis anterior muscle (TA) were tonically contracted. Muscle activity was measured using high-density electromyography, which allowed us to decompose the spiking activity of pools of motor units innervating the studied muscles. By analysing motor unit pool activity, we observed that none of the tested TACS conditions could consistently alter the spectral characteristics of the common neural inputs received by the muscles. These results suggest that 1mA-TACS over motor cortex given at frequencies in the beta band does not affect corticospinal beta entrainment.

**Highlights:**

- TACS is commonly used to entrain the communication between brain regions
- It is challenging to find direct evidence supporting TACS-driven neural entrainment
- Simulations show that motor neurons are sensitive to corticospinal beta entrainment
- Motor unit activity from human muscles does not support TACS-driven entrainment

## Introduction

In humans, transcranial alternating-current stimulation (TACS) has been used to entrain the outputs of the stimulated cortical areas and their synchronisation with other parts of the nervous system [1–4]. However, proof of TACS-driven entrainment is difficult to obtain since direct measurement of brain rhythms non-invasively during TACS is technically challenging [5–7]. At present, the extent to which TACS can induce changes in human brain activity remains unknown.

Animal studies suggest that TACS can acutely entrain cortical neuronal firing, especially when coupled with endogenous rhythmic brain activity [8,9]. A common form of TACS involves using stimuli at frequencies matching the beta oscillations (13-30Hz) that are observed in the motor cortex [10–12]. Corticospinal cells are involved in the generation of such motor cortical beta rhythms [13]. Therefore, if TACS can cause sufficiently strong levels of cortical beta entrainment, such effect is also expected to be apparent in the activity of corticospinal neurons. If this is the case then, given that the corticospinal tract can reliably transmit cortical beta rhythms to motor neurons (MNs) during tonic muscle contractions [14], it is expectable that TACS-induced corticospinal beta entrainment can be assessed by studying its distal effects on MNs [15–17]. This would provide a novel method to study TACS-driven neural entrainment based on its distal effects, which is especially attractive, as the large separation between cortical stimulation and muscle recordings will minimise contamination by the stimulus artifact.

We tested if TACS targeting the motor cortex can modulate the inputs received by a pool of MNs innervating a contracted muscle. This would provide an indirect evidence that TACS can entrain cortical activity. First, we used a computational model of a MN pool receiving different inputs to assess how reliably the common activity in the MN pool could inform about changes in corticospinal beta activity (considered a common input to MNs). Then, we ran an experiment in humans aimed to characterize TACS-induced changes in the firing activity of motor unit pools of upper- and lower-limb muscles during tonic contractions. Specifically, to test whether TACS was able to entrain cortical rhythms relayed through the corticospinal tract, we studied whether ongoing levels of common activity in the motor unit pools changed when TACS was delivered. TACS was given at frequencies in the beta band (21Hz), or at two control stimulation frequencies (7Hz and 40Hz) at which no corticomuscular interactions are normally found [11,18–20].

## Methods

This study comprises two parts. Part I simulates how the activity of a MN pool changes when the level of beta entrainment of corticospinal common projections to the MNs is modulated. Part II involves experiments using TACS and measuring muscle activity with high-density electromyography (HD-EMG) to measure TACS-driven changes in the neural drive to the muscle by analysing the spiking activity of pools of motor units.

### Part I - Simulation of a pool of MNs receiving common beta inputs

It has been previously shown that cortical oscillations are transmitted to the muscles through the fastest descending pathways [14]. This implies that, when simulating the activity of MN pools receiving cortical oscillatory inputs, one can use simplified models that only consider the fastest and most direct descending corticospinal projections to MNs. Here we used a computational model of a pool of MNs receiving a common input that simulated the summed contribution of corticospinal neurons, and independent inputs that were different for each MN (Fig. 1). Individual corticospinal inputs were simulated as spike trains with the times of the spikes randomly determined following Poisson distributions with a time-varying rate parameter lambda. Lambda was obtained by summing the average discharge rate of corticospinal neurons (set to 25 spikes/s [19,21]) with a sine wave at 21Hz with different amplitudes (simulating the modulation of the discharge rate of corticospinal neurons by beta inputs [14]). The net common input to MNs was the result of summing the spike trains of 100 corticospinal neurons, which simulated the contribution of the fastest corticospinal projections to the MN pool. The number of corticospinal neurons used was based on the relation between the estimated size of unitary excitatory postsynaptic potentials (EPSP) from corticomotoneuronal projections, and the size of compound EPSPs resulting from stimulating the pyramidal tract [19,22,23]. The independent inputs to MNs were modelled as white Gaussian noise, with mean equal to variance. The level of the mean was adjusted with reference to preliminary simulations to make MNs fire at an average discharge rate of ^~^11Hz [14]. To simulate the MN behaviour, we used a previously validated computational model [18,19,24]. The model was implemented in MATLAB (v-2020a, The Mathworks Inc., USA) and it simulated the firing activity of 177 MNs. The single MN models were conductance-based and they included somatic and dendritic compartments. The differential equations governing MN membrane potential were solved using the exponential integration scheme [25], with a time step of 0.2ms.

**Fig. 1.**
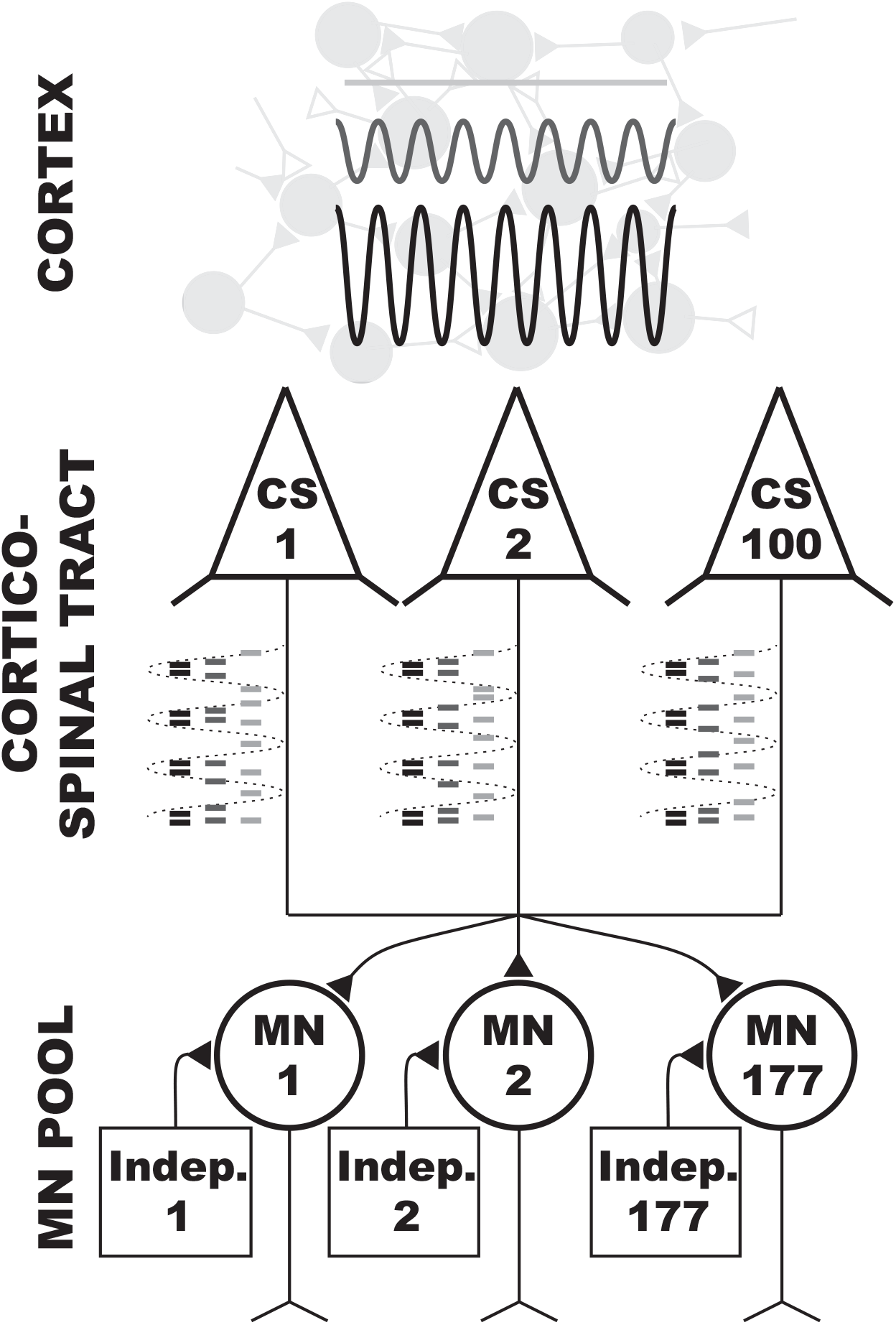
Schematic of the motor neuron (MN) model used in this study. A pool of 177 MNs receive independent inputs (different inputs to each MN) and a common input simulating the descending neural input from 100 corticospinal neurons (CS) with monosynaptic connection to MNs. The model is used to test how the estimated common inputs to MNs change as a function of the level of entrainment of the corticospinal tract with beta oscillations. Three levels of beta entrainment are exemplified using a grayscale.

To study how the activity of MNs changes due to changes in beta entrainment of corticospinal neurons, we ran a series of simulations in which we gradually increased the amplitude of the beta component modulating the discharge rates of the corticospinal neurons. To have a reference value for the amplitude of this beta component (referred to as ‘reference beta level’ from now on), we considered previous studies on primates looking at the typical values of coherence within the beta band between recordings of local field potentials (LFPs) in layer V of the primary motor cortex and the spike trains of corticospinal neurons [26]. Based on these works, peak beta coherences should be around 0.02. Therefore, we determined the amplitude of the beta signal (21Hz) modulating the discharge rate of corticospinal neurons that led to a coherence amplitude of 0.02 between the beta sinusoid and the spiking activity of the corticospinal neurons. The estimated reference beta level was 2.5 spikes/s, i.e., 10% of the baseline discharge rate of the corticospinal neurons. Based on this level, we ran 100 simulations of 121s each (with the first second discarded in the analysis) testing increasing levels of beta amplitude from zero (no modulation) to 4 times the reference beta level. These tests allowed us to model how reliably spiking activity of MNs can inform about changes in beta activity in the corticospinal tract.

We also used the MN model to estimate the expected minimum detectable effect size (MDES) of experiments in Part II. We ran 100 simulations in which the level of beta modulation of the corticospinal neurons was kept constant at the reference beta level. In this case, the simulated blocks were of 41s (with the first second discarded) to make them match with the analysed data in Part II. Results were then used to estimate the minimum level of change in the intramuscular coherence (the main outcome measure here, as described below) that should be detectable given the experimental conditions in Part II.

### Part II - Motor unit activity in contracted muscles during TACS

Here, we analysed how the common activity in pools of motor units innervating upper- and lower-limb muscles changed during TACS. For this purpose, we recruited ten healthy subjects (9 male; ages 22-40). All subjects provided written consent approved by the ethics committee of the University College London and in accordance with the Declaration of Helsinki (Ethics Application 10037/001). None of the participants had contraindications to TACS.

#### Experimental task

Recording sessions comprised two separate blocks in which we collected data from the right tibialis anterior (TA) and abductor digiti minimi (ADM) muscles during isometric contractions and while TACS was delivered over the motor cortex. Fig. 2A shows the position in which the arm and leg were held during the recordings. At the beginning of each block, the maximum voluntary isometric contraction (MVC) of the studied muscle was estimated. Each block consisted of four runs in which subjects departed from a relaxed position and followed a path on a screen by producing forces with the measured muscle. The target force path consisted of 1) a resting period (5s); 2) a ramp contraction period (5s) where force was linearly increased to reach a target level of 5% (ADM) or 10% (TA) of the MVC; and 3) 60s of steady contraction. The different contraction levels required for ADM and TA was based on the different characteristics of motor units in the two muscles, and they were meant to lead to the activation of large enough pools of units without causing fatigue [27].

**Fig. 2.**
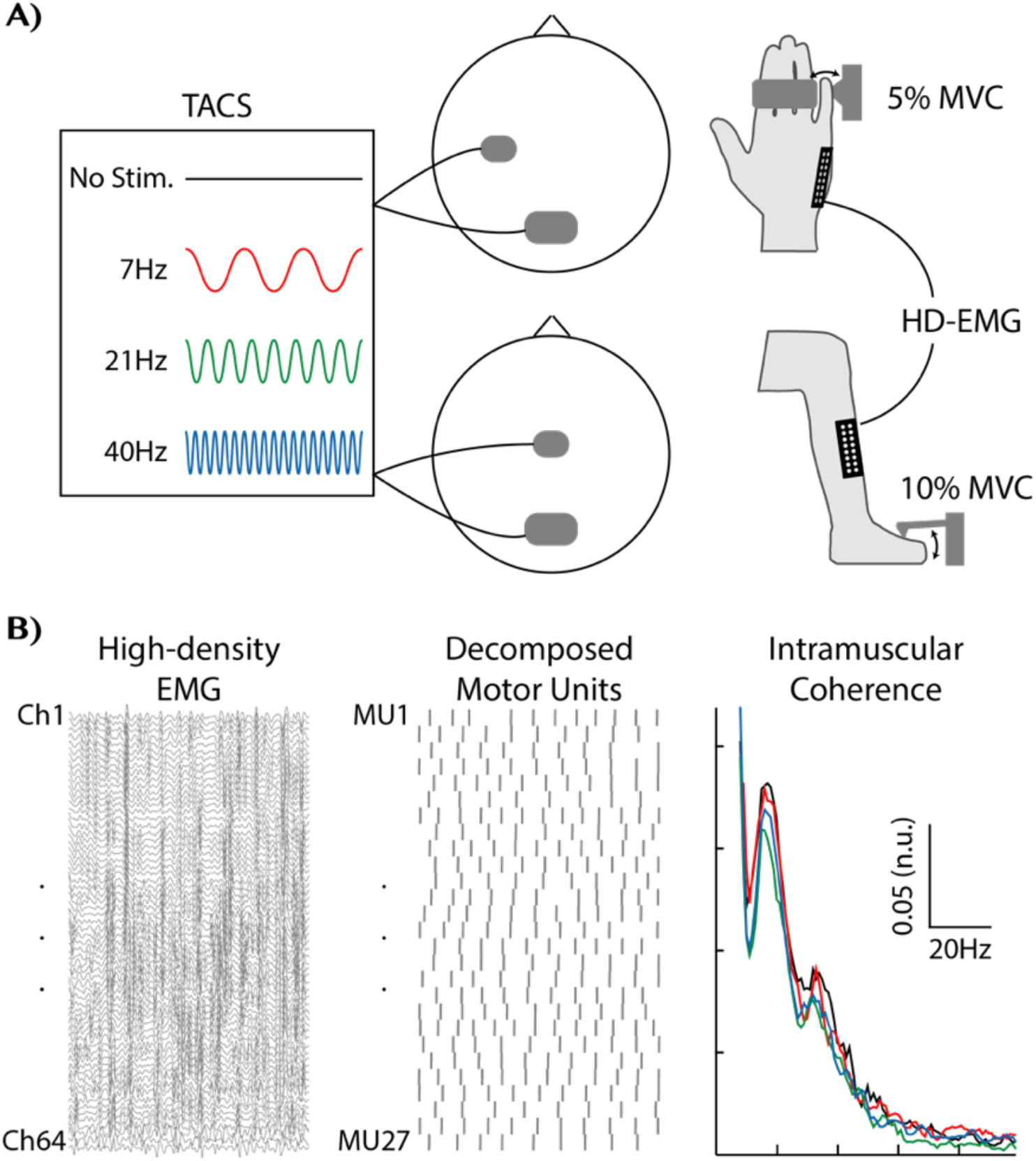
Experimental set up in Part II of the present study and recordings of the common inputs to a muscle using the intramuscular coherence function. (A) Four TACS conditions were delivered (No Stim. -black-; 7Hz TACS -red-; 21Hz TACS - green-; 40Hz TACS -blue-) either over the hand or leg cortical area while isometric steady contractions were produced either with the ADM or the TA. (B) HD-EMG recordings were used to extract information about spiking activity of motor units from the contracted muscles. Spiking activity of pools of motor units was used to estimate the common synaptic inputs to the motor neuron pools by computing the intramuscular coherence.

#### Stimulation

TACS with an amplitude of 1mA and at 7Hz, 21Hz and 40Hz was delivered through a pair of rubber electrodes (anode: 5×5cm^2^; cathode: 5×10cm^2^) adhered to the scalp using conductive paste (stimulator: DC-Stimulator plus, Neuroconn, Germany). The anode was placed either over C3 (ADM) or Cz (TA) positions on the scalp based on an EEG cap with a 10/20 layout (Fig. 2A). Pz was used for the cathode to avoid inducing phosphenes [12]. TACS was driven by a DAQ board (USB-6229, National Instruments, USA). In each run, the stimulation started with subjects at rest, and it continued during the entire duration of the isometric contraction of the muscle sustained for 60s, from which we analysed the last 40s. This was done to ensure that the TACS effects were stable during the analysed interval and to maximize the number of decoded motor units steadily firing [28].

#### Muscle recordings

HD-EMG grids with 64 contact points (13×5 matrices) and an inter-electrode distance of 4mm (ADM) or 8mm (TA) were placed centred around the innervation zone of the muscles after skin preparation (Fig. 2A). A bracelet around the distal part of the forearm was used as ground and an additional bracelet around the bony area of the wrist (ADM) or the ankle (TA) was used as the reference. EMG signals were band-pass filtered (20–500Hz) and sampled at 2048Hz (Quattrocento, OTBioelettronica, Italy).

#### Decoding of motor unit activity

HD-EMG signals were decomposed offline into motor unit spike trains using a validated blind source separation procedure [29] (Fig. 2B). The estimated motor unit spike trains were then visually inspected and processed following previously proposed guidelines [28,30]. From the decomposed motor units, only those active throughout the analysed intervals and with a pulse-to-noise ratio (PNR) over 30dB were kept for further analysis [29]. A minimum of 6 reliably identified units in all runs of a block was set as the criterion to keep recording blocks for subsequent statistical analysis. This was done to ensure that the pools of units considered could reliably characterize common inputs in different frequency bands [31] and it led to discarding the ADM blocks of 3 subjects. The resulting pools of motor units were used to characterize the common inputs to muscles.

### Analysis of common inputs from MNs activity

To study the common inputs to a MN pool within different frequency bands, we used the intramuscular coherence function (IMC) [32] (Fig. 2B). IMC was obtained by running 100 iterations of an algorithm that first randomly divided the pool of MNs considered into two sub-pools of equal size, and then calculated the spectral coherence between the cumulative spike trains obtained from the two sub-pools by summing the spiking activity of the MNs in each pool. Coherence was computed using 1-s segments and multi-taper method for spectral estimation (NW=2; Neurospec-2.11) [14,33].

In the case of the data obtained from the first simulations in Part I (increasing levels of beta), IMC was computed using sub-pools of increasing sizes from 1 to 88 MNs and the average IMC levels within the 20-22 Hz band were determined for each level of beta input simulated.

For the data from the second set of simulations in Part I, IMC was computed using sub-pools of 7 MNs to approximate the average number of units reliably identified in Part II (13.4 units decomposed on average; see results). The mean (*μ*) and variance (*σ^2^*) of the average IMC in the 20-22Hz band across the 100 simulations were obtained and used to estimate the MDES as follows:

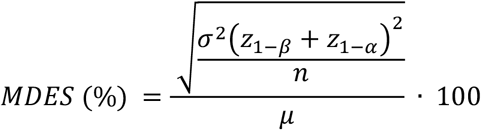

where *n* is the sample size (17 recordings considered in the main results in Part II), *α* and *β* are the probabilities of type I/II errors, and *z* refers to the critical Z value [34]. *α* and *β* were set to 0.05 and 0.2.

In Part II, the number of motor units used to estimate the IMC was determined for each subject by considering the block with the lowest number of units decomposed. In this case, we assessed if IMC amplitudes changed at the frequencies at which TACS was delivered. Therefore, we calculated the average IMC amplitudes within three frequency ranges: 4-13Hz; 13-30Hz; 30-50Hz and used these results to run statistical tests (the results section is based on data using these frequency bands, but we also include in the Supplementary files results of equivalent analyses using narrow windows of 2-Hz around the stimulus frequencies). Mixed modelling was used to determine the influence of TACS on IMC. Fixed factors included in the model were TACS stimulus (STIM; i.e. No Stim., 7Hz, 21Hz, 40Hz), frequency band of analysis (FREQ), and the interaction between the two (STIM x FREQ). Subject was included as a random factor. To test our specific hypotheses regarding the influence of TACS at 21Hz on beta coherence, we ran pairwise comparisons contrasting the TACS protocols. Results from the two muscles studied were merged for this analysis (we also include results of tests using muscle as a factor and muscle-specific tests in the Supplementary files). Assumptions of normality and homoscedasticity of the residuals were assessed visually using q-q plots and fitted-versus residual-value plots. The lmer package (Bates et al., 2014) in R (R Core Team, 2019) was used.

## Results

### Part I – MNs can reliably inform about changes in beta inputs to corticospinal neurons

Fig. 3 shows how common inputs to the simulated MN pool are measured by the IMC as the number of MNs considered increases. These results are for the case in which a reference beta level (estimated, as described in Methods, based on [26]) is used to modulate the activity of corticospinal neurons. Although only one sinusoidal component (the beta input at 21 Hz) modulates the common inputs, since corticospinal inputs to MNs are spike trains following Poisson distributions, their spectral contents cover a wide range of frequencies [35]. Therefore, variable levels of common inputs (*i.e.*, non-zero coherence levels) are observed in the IMC at frequencies outside the bandwidth of the beta input (Fig. 3A).

**Figure 3.**
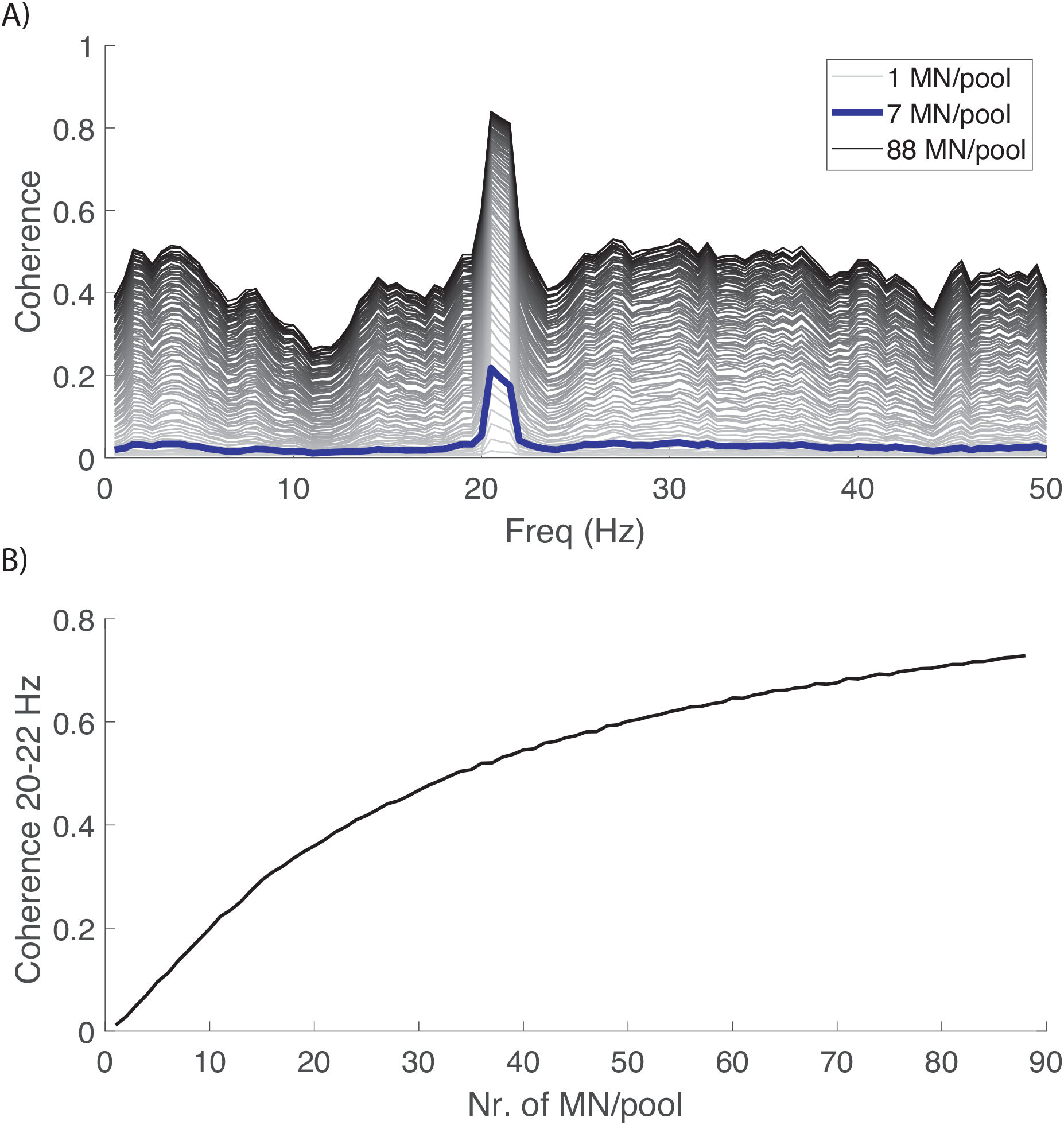
Intramuscular coherence estimated for the case in which a reference beta level is used to modulate corticospinal neurons firing activity. Results obtained using different number of motor neurons per pool (MN/pool) to estimate intramuscular coherence. A) Intramuscular coherence in the 0-50 Hz range (from light-grey to black, traces represent intramuscular coherence estimates using from 1 to 88 MNs/pool; the case in which 7 MNs per pool are considered is highlighted in blue); B) Average intramuscular coherence in the 20-22 Hz band as a function of the number of MNs used to estimate it.

As expected, the amplitudes of the IMC at the frequencies of the beta inputs to the MN pool increase when the size of the pools of MNs considered increases (Fig. 3A) [36]. However, when large pools of MNs are used to estimate the IMC, even very small common inputs outside the beta band (that may be spurious and due to chance) can be strongly enhanced by the IMC (as observed in the offset level present in the darkest traces in Fig. 3A). This may affect the characterization of actual common inputs (like beta inputs in these simulations), as the range between chance-related IMC levels and the maximum possible coherence of 1 shrinks. Interestingly, the computation of IMC from small-to-medium pools (*i.e.*, 10-30 MNs) results in close to zero coherence levels at frequencies outside the common beta inputs, while coherence at the beta input frequency vary with the input. This is the case, for example, when the IMC is computed from pairs of pools of 7 MNs each (blue trace, Fig. 3A), which matches with the average number of MNs that we could reliably decompose from real muscle recordings in Part II.

Before analysing how IMC changes with changes in the beta input to corticospinal neurons in our simulations, it is important to assess if the reference beta amplitude used (based on intracortical recordings in primates; see methods) produces IMC levels in the beta range similar to those observed experimentally. This should be expected if the only or most dominant common beta input to the MN pool resulted from a single -corticospinal-source. This is the case here: the IMC amplitude at 21Hz increases with the number of MNs used to estimate it, showing an initial steep increase followed by a slower ramp trending towards 1 (Fig. 3B). Based on this graph, when the number of MNs used approximates what is typically decoded in human experiments (*i.e.,* estimates of IMC based on 5-15 MNs/pool [28]), the IMC beta level is approximately 0.1-0.3. This is in line with human recordings during steady contractions [37,38].

The IMC is highly sensitive to changes in the beta modulation of corticospinal neurons. Fig. 4 shows how IMC changes with increasing amplitudes of the beta signal modulating corticospinal neurons and when either 1, 7 or 88 MNs per pool are used to estimate the IMC (Fig. 4A-C). The amplitude of IMC around 21Hz (frequency of the common beta input) follows the increases in the amplitude of the beta inputs. The number of MNs considered influences how changes in beta inputs are reflected in the IMC. Changes in the IMC when 1 or 88 MNs/pool are considered are constrained to coherences between 0 and ^~^0.4 and ^~^0.4 and 1. When the IMC is estimated using 7 MNs/pool, the beta IMC resulting from different levels of beta inputs range between 0 and ^~^0.9.

**Figure 4.**
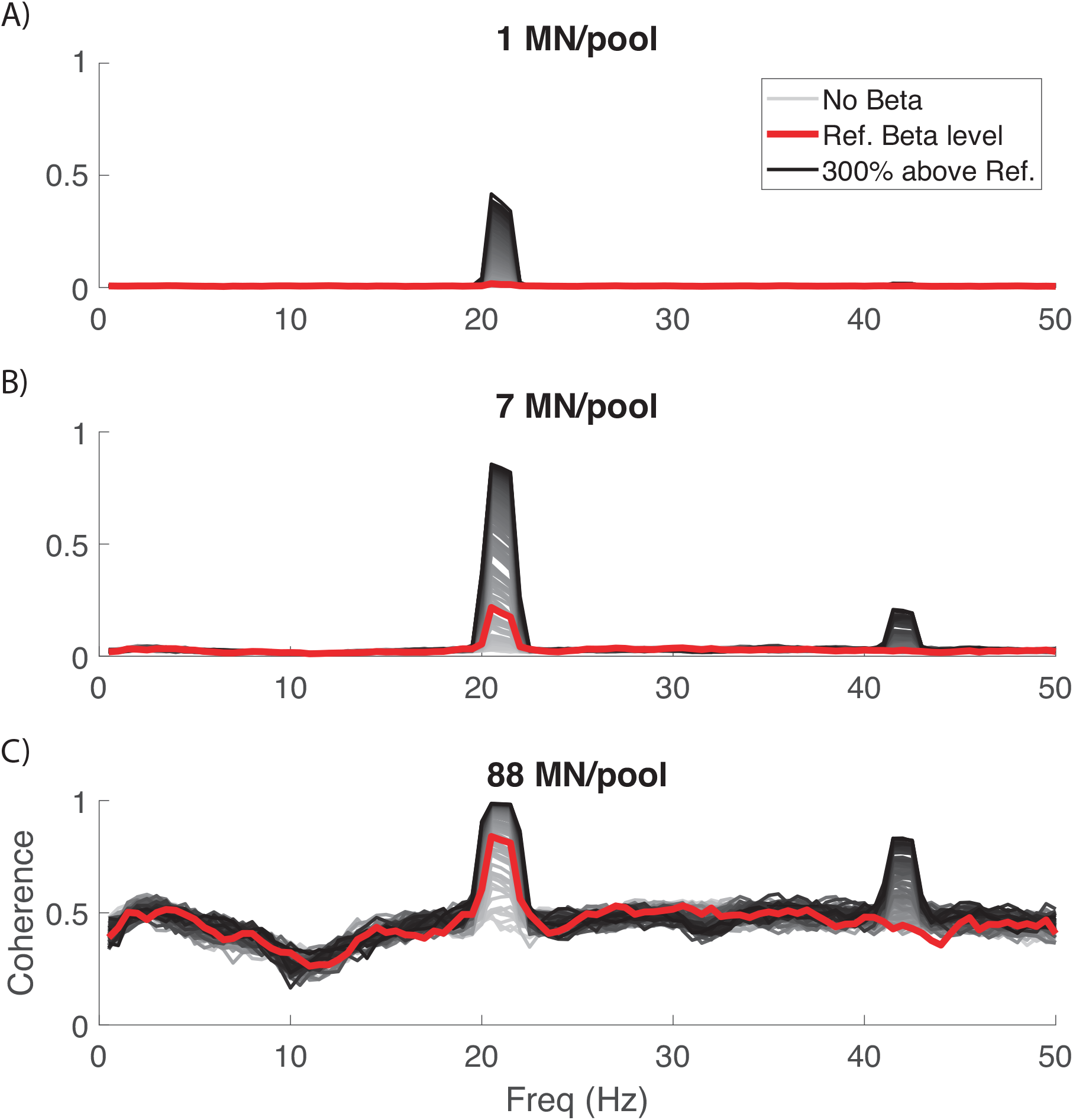
**Intramuscular coherence** with different amplitudes of the beta signal modulating pyramidal tract neurons projecting onto a simulated pool of motor neurons (MNs). A, B and C panels present results for the cases in which 1, 7 or 88 MNs/pool are considered to estimate the intramuscular coherence. From light-grey to black, traces represent the different levels of beta simulated ranging from no beta modulation to 300% increase in beta relative to the estimated reference beta level. The highlighted red trace represents the intramuscular coherence when the beta signal modulating inputs has the estimated reference beta level.

IMC changes with the amplitude of the beta signal modulating corticospinal activity. This is shown in Fig. 5 both in absolute terms (Fig. 5A) and in terms of changes in IMC relative to levels observed when the reference beta level is used as input (Fig. 5B). These results suggest a nearly linear relationship between moderate variations in the amplitude of the beta modulation relative to the estimated reference beta level, and changes in the IMC at beta frequencies. As a reference, these results suggest that changes of 8-12% in the amplitude of the beta signal modulating corticospinal activity (relative to the reference levels) should result in changes of approximately 0.015-0.030 in the IMC when 14 MNs (7 MNs/pool) are considered.

**Figure 5.**
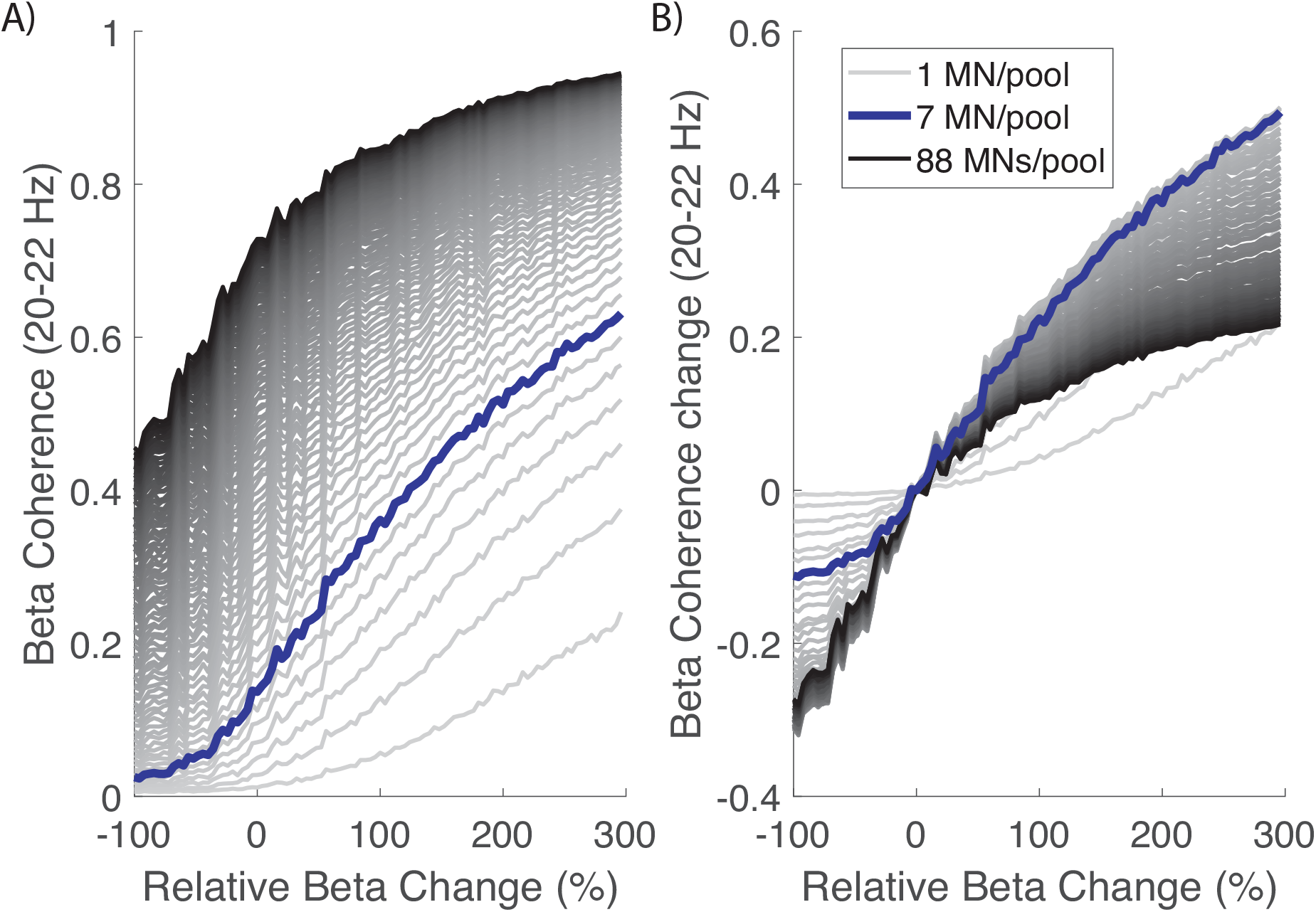
**Intramuscular coherence in the 20-22 Hz band** as a function of changes in the amplitude of the beta signal modulating the firing activity of corticospinal neurons. A) Absolute intramuscular coherence amplitudes as a function of changes in beta inputs; B) Changes in intramuscular coherence amplitudes relative to coherence when using the reference beta level.

Finally, an MDES of 7% was obtained based on the results from the second set of simulations in Part I. This implies that the experimental conditions in Part II are expected to be powered to detect TACS-driven changes in IMC greater than 7% relative to baseline.

### Part II – Estimated common inputs to muscles in humans do not change during TACS

Across subjects and muscles, 13±4 motor units were identified (range 6-22; 9.7±2.4 ADM; 15.2±4.1 TA). The average discharge rate of the motor units during steady contractions was 11.9±2.1 spikes/s. Paired t-tests run between all tested conditions showed no significant effect of TACS on average forces (*p*>.3 in all paired comparisons).

Fig. 6 summarizes individual and group IMC results. We did not find differences in the IMC between the tested TACS conditions. Specifically, results from the model examining IMC changes indicated no effect of STIM (ß=-3.74e-04, SE=8.24e-04, p=0.65; Table 1), and no significant STIMxFREQ interaction (ß=2.65e-06, SE=3.12e-05, p=0.93). Paired comparisons showed a difference between blocks with no TACS and with TACS given at 21Hz on the IMC levels in the beta band (p=0.027; see Supp. Table 1), suggesting that the amplitude of the IMC in the beta band decreased during 21Hz-TACS. However, this significance did not survive post-hoc correction for multiple comparisons.

**Fig. 6.**
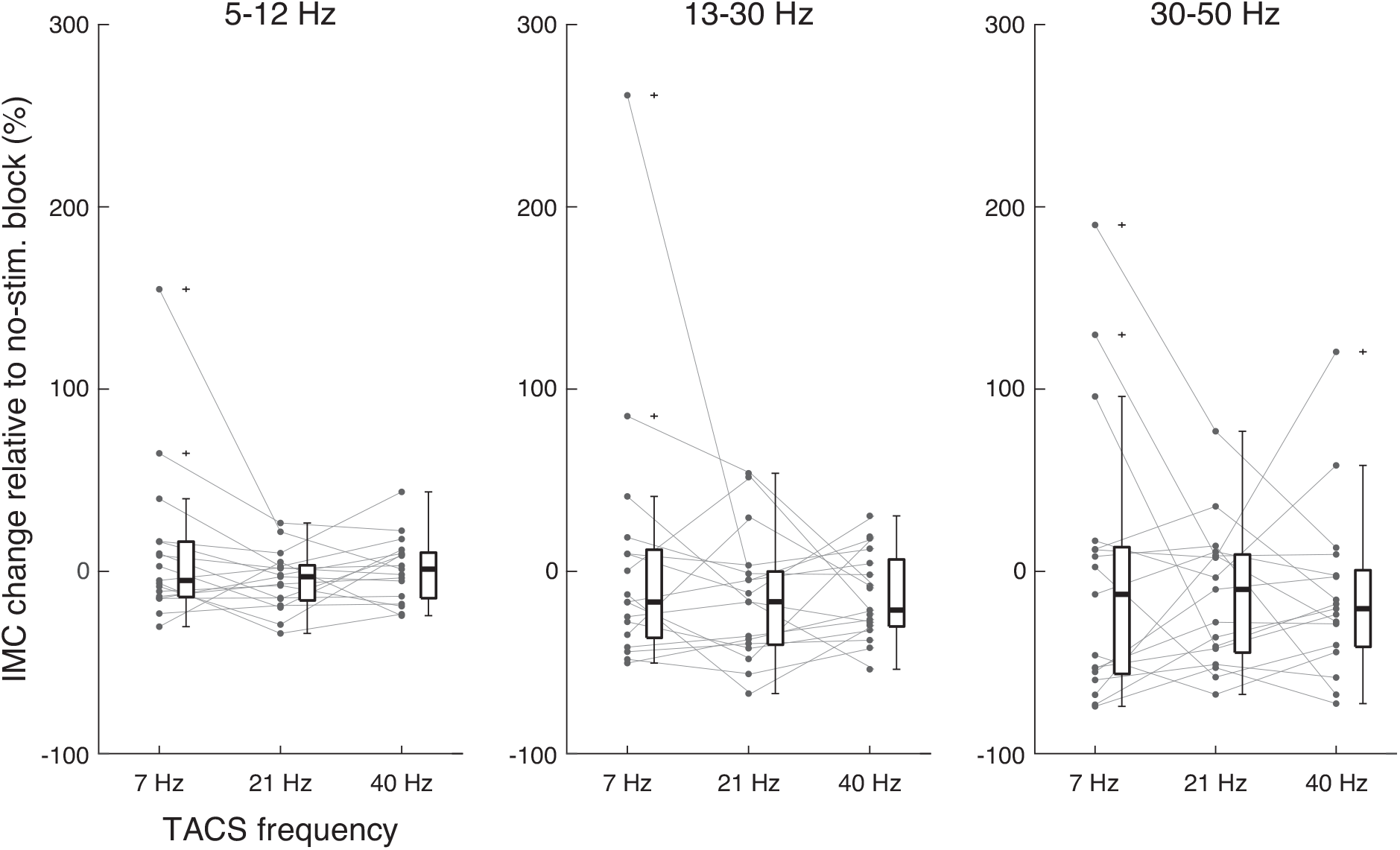
Changes in intramuscular coherence during TACS. Intramuscular coherence changes (relative to No Stim blocks) within three frequency bands of interest and for the three TACS conditions tested: TACS given at 7Hz, 21Hz or 40Hz). Results with TA and ADM muscles are merged. Individual results are represented by the connected dots. Boxplot are included to compare results between groups.

**Table 1:** Results with the main model used in the study to examine intramuscular coherence changes due to TACS. There was no significant influence of TACS on the IMC, and there was no interaction between the TACS protocol used (STIM) and the frequency band considered (FREQ).

| Model Parameter | Estimate | SE | 95% Confidence Interval | p-value |
| --- | --- | --- | --- | --- |
| Intercept | 3.13e-01 | 2.96e-02 | 2.51e-01 to 3.76e-01 | <0.001* |
| STIM | -4.13e-04 | 8.25e-04 | -2.04e-03 to 1.21e-03 | 0.617 |
| FREQ | -6.66e-03 | 7.15e-04 | -8.07e-03 to -5.25e-03 | <0.001* |
| STIM x FREQ | 3.64e-06 | 3.13e-05 | -5.80e-05 to 6.53e-05 | 0.907 |

Analogous tests to the main one presented above were also performed considering IMC levels in narrow bands of 2Hz around the frequencies used for stimulation (Supp. Table 2). Additionally, tests were also performed using muscle-specific data (Supp. Tables 3-4) and using muscle as a random factor in the analysis (Supp. Table 5). The main results did not change in any of these cases as we did not find significant effects of factor STIM or of STIM x FREQ interaction found.

## Discussion

Finding direct *in vivo* evidence of the effects of TACS on ongoing neural activity in an undamaged human brain is challenging due to technical limitations in existing brain recording technologies [39]. Here, we propose a way to infer TACS-driven corticospinal entrainment by assessing the distal effects that the stimulation has on alpha motor neurons (MNs) innervating muscles. The experiments run to test this methodology lead to two contrasting results: while simulations indicate that information from pools of MNs in a muscle can reliably inform about changes in corticospinal entrainment, results from human experiments show that TACS over the motor cortex does not change the spectral properties of the common inputs received by pools of MNs in upper- and lower-limb muscles. Considering the involvement of corticospinal neurons in the generation and propagation of beta rhythms observed in the motor cortex [13], our results also suggest that TACS, with the intensity and montage used here, does not have a strong effect on motor cortical neural activity.

Recent studies testing TACS in primates have suggested that the activity of some cortical neurons can be entrained by TACS [8]. Based on our simulations, the assessment of common inputs to MNs in activated muscles should be able to inform about relatively small levels of entrainment in the corticospinal tract. However, results from our human experiments suggest that common inputs remain largely unchanged during TACS. In fact, stimulation with beta frequencies not only did not increase levels of common beta inputs to MNs, but it showed a trend towards the opposite direction. This lack of evidence for rhythmic entrainment may be interpreted in different ways. First, our results may indicate that TACS delivered with standard intensities in human experiments [12] is unable to entrain neural activity of, at least, MNs and the pyramidal cells with monosynaptic connections with them. Future studies should be performed to assess if TACS given using higher currents can produce observable effects [12]. A second explanation for our results may be that entrainment using standard TACS intensities is only possible when the stimulated brain areas are in a dynamic phase (*i.e.,* transitioning between two states), since during these less stable neural states external stimuli appear to be more capable of producing changes in the brain [40,41]. This would explain why we could not find any evidence for corticospinal entrainment here while previous works (relying on scalp electrical signals during TACS) found significant TACS-driven corticomuscular beta entrainment during periods preceding movements [42]. Finally, it may be the case that TACS is only capable of entraining corticospinal activity when falling in-phase with endogenously generated beta activity, while the induced effects may be strongly reduced when stimulation presents random phases relative to cortical beta activity. Future studies may be able to test this possibility by driving TACS in closed-loop with estimates of ongoing cortical beta activity [43].

To simulate how changes in corticospinal beta entrainment affect MNs’ activity, we used a simplified model of a MN pool receiving common inputs from a relatively small pool of corticospinal cells representing the fastest brain-muscle pathways [44–46]. This is motivated by previous works showing that brain oscillations projected to muscles travel through the fastest pathways [14]. Interestingly, when the model is run using experimentally observed levels of corticospinal entrainment to cortical beta rhythms in primates [26], the levels of intramuscular beta coherence resemble those found in human experiments [37,38]. This supports the suitability of the model and allows us to use a realistic reference value to study the effects of corticospinal beta entrainment. Based on this, the model leads to two predictions about the effects of changes in corticospinal beta entrainment on MNs’ activity. First, it shows that there is a nearly linear relationship between small deviations from the used beta reference level in the corticospinal inputs and the observed changes in the intramuscular coherence function. Second, it shows that relatively small pools of MNs (10-30 MNs) can readily provide an optimal description of changes in the beta common inputs. This is a relevant outcome to validate the results obtained in real experiments and based on information from limited pools of motor units due to the limitations in extracting information from non-invasive recordings of muscle activity [29,47].

Several limitations should be considered here. First, we did not use subject-specific current flow modelling and, therefore, induced currents may have been non-equally effective across subjects [48]. Since the stimulus intensities were similar to those in previous works [11,49], we estimate that our group results faithfully represent the effects of TACS on MNs. It is also not possible to determine the relevance of non-cortical common inputs to a MN pool, which may affect the strength with which cortical inputs are seen in pools of motor units. Since our simulations using beta modulation levels based on primate data led to beta IMC levels similar to those found in real experiments, we do not expect that there exist other relevant non-cortical beta sources to the muscles. Finally, TACS was delivered at fixed frequencies. Future studies should assess if adapting TACS to subject-specific beta frequencies leads to different outcomes [9]. Since we did not find clear evidence for entrainment on a subject-by-subject level, we do not expect that this factor has a major impact on our conclusions.

## Conclusion

We aimed to motivate the study of TACS-driven neural entrainment by looking at the distal effects on MNs. This was supported by realistic simulations suggesting that common inputs to MNs should be sensitive to changes in corticospinal entrainment. However, our experimental results indicate that TACS could not alter MN activity, which suggests that TACS-driven motor cortical entrainment may not be easy to achieve.

## Acknowledgements

We thank Paul Hammond for building the experimental platform used to record forces generated by the tibialis anterior muscle. This work was supported by the European Research Council Synergy Grant Natural BionicS (Contract #810346). JI received the support from “la Caixa” Foundation (ID 100010434; fellowship code LCF/BQ/PI21/11830018).

## CRediT authorship contribution statement

JI: Conceptualization, Methodology, Software, Formal analysis, Investigation, Writing - original draft, Writing - review& editing, Visualization. BZ: Investigation, Methodology, Writing - review & editing. KB: Investigation, Methodology, Writing - original draft, Writing - review & editing. LR: Methodology, Writing - review & editing, Visualization. ADV: Conceptualization, Investigation, Writing - review & editing. DS: Conceptualization, Writing - review & editing. CAV: Methodology, Formal Analysis. JR, DF, SNB: Conceptualization, Resources, Writing - review & editing, Funding acquisition.

## Supplementary files

**Supp. Table 1:** P-values of the pairwise comparisons between IMC amplitudes within the beta band (13-30Hz) observed in the different TACS conditions tested. Results before correction applied for multiple comparisons. * for p-values < 0.05.

| TACS condition (STIM) | No TACS | 7Hz | 21Hz |
| --- | --- | --- | --- |
| 7Hz | 0.987 |  |  |
| 21Hz | 0.027* | 0.126 |  |
| 40Hz | 0.204 | 0.334 | 0.517 |

**Supp. Table 2:** Model examining IMC amplitudes within 2-Hz bands around the stimulus frequencies (i.e., 7Hz, 21Hz, 40Hz). There was no significant influence of the TACS condition (STIM) on IMC, and there was no interaction between STIM and the analysed frequency band (FREQ). * for p-values < 0.05.

| Model Parameter | Estimate | SE | 95% Confidence Interval | p-value |
| --- | --- | --- | --- | --- |
| Intercept | 3.91e-01 | 4.43e-02 | 3.02e-01 to 4.80e-01 | <0.001* |
| STIM | -2.43e-04 | 1.53e-03 | -3.25e-03 to 2.77e-03 | 0.874 |
| FREQ | -8.72e-03 | 1.32e-03 | -1.13e-02 to -6.12e-03 | <0.001* |
| STIM x FREQ | -6.22e-06 | 5.78e-05 | -1.20e-04 to 1.08e-04 | 0.914 |

**Supp. Table 3:** Model examining IMC amplitudes considering only data from the ADM muscle. Similarly to the results in the main analysis, no significant effects were found that would indicate corticospinal entrainment at the TACS frequency. * for p-values < 0.05.

| Model Parameter | Estimate | SE | 95% Confidence Interval | p-value |
| --- | --- | --- | --- | --- |
| Intercept | 3.14e-01 | 3.00e-02 | 2.52e-01 to 3.76e-01 | <0.001* |
| STIM | -4.13e-04 | 8.25e-04 | -2.04e-03 to 1.21e-03 | 0.617 |
| FREQ | -6.66e-03 | 7.15e-04 | -8.08e-03 to -5.25e-03 | <0.001* |
| STIM x FREQ | 3.64e-06 | 3.13e-05 | -5.80e-05 to 6.53e-05 | 0.907 |

**Supp. Table 4:** Model examining IMC amplitudes considering only data from the TA muscle. Similarly to the results in the main analysis, no significant effects were found that would indicate corticospinal entrainment at the TACS frequency. * for p-values < 0.05.

| Model Parameter | Estimate | SE | 95% Confidence Interval | p-value |
| --- | --- | --- | --- | --- |
| Intercept | 2.73e-01 | 3.16e-02 | 2.08e-01 to 3.38e-01 | <0.001* |
| STIM | -3.84e-04 | 8.78e-04 | -2.12e-03 to 1.36e-03 | 0.662 |
| FREQ | -4.94e-03 | 7.60e-04 | -6.44e-03 to -3.43e-03 | <0.001* |
| STIM x FREQ | -6.21e-06 | 3.33e-05 | -7.21e-05 to 5.97e-05 | 0.852 |

**Supp. Table 5:** Model examining IMC amplitudes considering muscle as a random factor. Although there were differences between the IMC levels in the muscles tested (mainly due to the different amount of units decomposed in each case), the main results in our study were not altered when using muscle as a random factor. * for p-values < 0.05.

| Model Parameter | Estimate | SE | 95% Confidence Interval | p-value |
| --- | --- | --- | --- | --- |
| Intercept | 4.74e-01 | 6.60e-02 | 3.44e-01 to 6.04e-01 | <0.001* |
| STIM | -5.25e-04 | 2.67e-03 | -5.79e-03 to 4.74e-03 | 0.844 |
| FREQ | -1.33e-02 | 2.31e-03 | -1.79e-02 to -8.74e-03 | <0.001* |
| MUSCLE | -1.00e-01 | 3.70e-02 | -1.73e-01 to 2.74e-02 | 0.007* |
| STIM x FREQ | 4.16e-05 | 1.01e-04 | -1.58e-03 to 2.41e-04 | 0.681 |
| STIM x MUSCLE | 7.05e-05 | 1.61e-03 | -3.09e-03 to 3.24e-03 | 0.965 |
| FREQ x MUSCLE | 4.20e-03 | 1.39e-03 | 1.45e-03 to 6.94e-03 | 0.002* |
| STIM x FREQ x MUSCLE | -2.39e-05 | 6.08e-05 | -1.44e-03 to 9.60e-05 | 0.694 |

